# *NF1* deficiency correlates with estrogen receptor signaling and diminished survival in breast cancer

**DOI:** 10.1101/272815

**Authors:** Patrick S. Dischinger, Elizabeth A. Tovar, Curt J. Essenburg, Zachary B. Madaj, Eve Gardner, Megan Callaghan, Ashley N. Turner, Anil K. Challa, Tristan Kempston, Bryn Eagleson, Robert A. Kesterson, Roderick T. Bronson, Megan J. Bowman, Carrie R. Graveel, Matthew R. Steensma

**Affiliations:** Center for Cancer and Cell Biology, Van Andel Research Institute, Grand Rapids, MI, USA; Bioinformatics & Biostatistics Core, Van Andel Research Institute, Grand Rapids, MI, USA; Department of Genetics, The University of Alabama at Birmingham, Birmingham, AL, USA; Vivarium and Transgenics Core, Van Andel Research Institute, Grand Rapids, MI, USA; Rodent Histopathology Core Dana Farber/Harvard Cancer Center, Harvard Medical School, Boston, MA, USA; Helen DeVos Children’s Hospital, Spectrum Health System, Grand Rapids, MI, USA; Michigan State University College of Human Medicine, Grand Rapids, MI, USA

## Abstract

The key negative regulatory gene of the RAS pathway, *NF1*, is mutated or deleted in numerous cancer types and is associated with increased cancer risk and drug resistance. Even though women with neurofibromatosis (germline *NF1* mutations) have a substantially increased breast cancer risk at a young age and *NF1* is commonly mutated in sporadic breast cancers, we have a limited understanding of the role of *NF1* in breast cancer. Much of our understanding of the mechanisms underlying the functional loss of *NF1* comes from mouse models that do not completely recapitulate the phenotypes of human *NF1*. We utilized CRISPR-Cas9 gene editing to create *Nf1* rat models to evaluate the effect of *Nf1* deficiency on tumorigenesis. The resulting *Nf1* indels induced highly penetrant, aggressive mammary adenocarcinomas that express estrogen receptor and progesterone receptor. We identified distinct *Nf1* isoforms that were altered during tumorigenesis.

To evaluate *NF1* in human breast cancer, we analyzed genomic changes in a breast cancer dataset of 2,000 clinically annotated breast cancers. We found *NF1* shallow deletions in 25% of sporadic breast cancers, which correlated with poor clinical outcome. To identify biological networks impacted by *NF1* deficiency, we constructed gene co-expression networks using weighted gene correlation network analysis (WGCNA) and identified a network connected to *ESR1* (estrogen receptor). Moreover, *NF1*-deficient cancers correlated with established RAS activation signatures. Estrogen-dependence was verified by estrogen-ablation in *Nf1* rats where rapid tumor regression was observed. These results demonstrated the significant role NF1 plays in both NF1-related breast cancer and sporadic breast cancer.

## Introduction

Deregulated RAS signaling promotes several “hallmarks of cancer”, such as sustained proliferation, invasion and metastasis, and angiogenesis.^1^ The importance of RAS deregulation in cancer is demonstrated by the fact that *KRAS* is the most commonly mutated oncogene and occurs at a high frequency in lung (17%), pancreatic (57%), and colon (33%) cancers.^2^ Even though KRAS is mutated in only 4% of sporadic breast cancers (*HRAS* = 1%; *NRAS*=2%)^2^, the RAS/ERK pathway is hyperactivated in approximately 50% of breast cancers.^3–5^ This discrepancy suggests that there is another mechanism underlying RAS activation, besides mutation, in human breast cancers. The key negative regulatory gene of the RAS pathway, *NF1*, is mutated or deleted in a wide range of cancers and is increasingly recognized as a significant cancer driver. The *NF1* gene encodes neurofibromin, a GTPase-activating protein that regulates RAS (including HRAS, NRAS, and KRAS) and loss of neurofibromin function results in hyperactivated RAS.^6^ Exons 20-27 encode the GAP-related domain (GRD), which is the RAS regulatory domain of human neurofibromin.^7^ Mutations or deletions within the GRD result in decreased NF1 functionality and dysregulated RAS signaling driven through the RAS–RAF–MEK–ERK cascade.^8^ Neurofibromatosis type 1 (NF1) is caused by germline mutations in the *NF1* gene and is the most common single-gene disorder affecting 1 in 3,000 live births.^9,10^ The majority of NF patients develop benign cutaneous neurofibromas and may also develop peripheral nerve tumors, neurocognitive disorders, and bone stigmata (tibial dysplasia, scoliosis, and osteoporosis). Importantly, NF patients have an increased risk of developing several adult cancers including breast, ovarian, liver, lung, bone, thyroid, and gastrointestinal cancers.^11^ These diverse clinical manifestations reveal the impact of loss of NF1 function and dysregulated RAS in numerous tissue types.

In addition to germline mutations, *NF1* mutations and deletions commonly occur in sporadic cancers and are associated with increased cancer risk and drug resistance. *NF1* is the third most prevalent mutated or deleted gene in glioblastoma,^12^ fourth most mutated gene in ovarian cancer,^13^ and the second most common mutated tumor suppressor in lung adenocarcinoma.^14^ More recently, clinical evidence has mounted demonstrating that women with NF1 have a significantly increased breast cancer risk. A risk analysis of 3,672 NF patients found that women with NF1 have an increased relative risk of developing breast cancer in their younger years compared to the general population (relative risk was 6.5 at 30-39 years, 4.4 at 40-49 years, and 2.6 at 50-59 years).^15^ Another key study of 1,404 NF patients identified an unequivocal increased risk for breast cancer with standardized incidence ratios of 11.1 (95% CI, 5.56 to 19.5) for breast cancer in women with NF1 age < 40 years and the overall breast cancer mortality ratio was 5.20 (95% CI, 2.38 to 9.88).^16^ This study, in addition to several others, has established the increased breast cancer risk and associated poor outcome in patients with neurofibromatosis.^17^ Comprehensive genomic analyses of sporadic breast cancers revealed that NF1 is commonly mutated and may be an important driver in sporadic breast cancer.^18,19^ A study using the *Mcm4*^*Chaos3*^ mouse model of chromosomal instability identified *Nf1* deletions in almost all of the *Mcm4*^*Chaos3*^ mammary adenocarcinomas.^20^ These findings suggest that NF1 is a critical tumor suppressor and potential driver of breast tumorigenesis.

Much of our understanding of the mechanisms underlying the functional loss of NF1 and tumorigenesis come from studies of genetically engineered mouse models.^21–24^ These models have been valuable in defining how NF1 loss and deregulated RAS/MAPK signaling promote tumorigenesis. However, mouse models have general limitations, particularly with respect to reproducing human pharmacokinetics and recreating putative interactions between genetic/environmental factors and induced gene deficiency.^25^ It is well established that mice with germline heterozygous *Nf1* mutations do not spontaneously express key aspects of the human phenotype and require additional crosses into other germline mutants such as *Tp53*, or cell-specific conditional *Nf1* mutation, to elicit more representative phenotypes.^22,26–30^Prior to the demonstration of CRISPR-Cas9 gene editing, manipulation of the rat genome was far more difficult than the mouse. The rat offers significant advantages over the mouse because of its larger size, more representative physiology to human disease, higher degree of cognition and memory, and ease of use in pharmaceutical studies.^31,32^ In our study we utilized CRISPR-Cas9 gene editing capabilities to create several rat models of *Nf1* deletions in order to evaluate the effect of *Nf1* deficiency on tumorigenesis. The resulting *Nf1* indels induced highly penetrant, aggressive mammary adenocarcinomas in multiple rat founder lines. Moreover, both male and female *Nf1*-mutant animals developed mammary adenocarcinomas.

To evaluate the impact of *NF1* in sporadic breast cancer we analyzed genomic changes in a large breast cancer dataset composed of more than 2,000 clinically annotated breast cancers. We found that *NF1* shallow deletions are present in 25% of sporadic breast cancers and correlated with poor clinical outcome.^33^ To identify biological networks impacted by *NF1* deficiency impacts we constructed gene co-expression networks using weighted gene correlation network analysis (WGCNA) and identified co-expression networks. A module associated with *NF1* shallow deletion contained several genes that are considerably important in both ER+ breast cancers and endocrine resistance, including *ESR1* and *FOXA1*. Unsupervised hierarchical clustering revealed that breast cancers with *NF1* shallow deletions form a distinct cluster that correlate with ER-negative breast cancer and RAS activation. We validated estrogen dependence of *Nf1*-deficient breast cancers by ablating mammary tumors in our *Nf1* rat model by ovarectomy. These results demonstrated the significant role *NF1* plays in both NF1-related breast cancer and sporadic breast cancer. Moreover, this novel *Nf1* rat model is invaluable for interrogating the role of NF1, estrogen-dependent breast cancer, and deregulated RAS signaling in sporadic and inherited breast cancer.

## Results

### Establishing *Nf1* rat models using CRISPR −Cas9 nucleases

To investigate the effects of altered *Nf1* function in rat, we used two unique sgRNAs to target the GRD region in exon 20 and disrupt neurofibromin function (Figure 1A). Two unique CRISPR/sgRNAs were synthesized and co-injected with Cas9 mRNA into one-cell-stage Sprague-Dawley rat embryos, which were then transferred to pseudopregnant females. From two rounds of injections, 19 pups were born. Genotyping was performed by amplifying a 452 bp fragment encompassing the sgRNAs target sites from DNA isolated from tail biopsies, followed by a heteroduplex mobility assay (HMA).^34^ PCR amplicons with distinct HMA profiles were cloned into a plasmid vector and plasmids from multiple individual colonies were sequenced to identify the genetic lesions resulting from non-homologous end joining (NHEJ). Based on HMA profiles, indels were identified in 18 of 19 pups (Supplementary Tables 1 and 2). Multiple alleles were present in the majority of the G0 animals and were confirmed by Sanger sequencing, which revealed 34 mutant alleles, including 25 unique mutant alleles (Supplementary Table 2). Of the 34 mutations observed, 25 (73.5%) were frameshift mutations in both the 5’ and 3’ CRISPR target sites; 14 mutations (41.2%) were deletions ranging from 54-63 bp, spanning both the target sites. Interestingly, all the large deletions were in-frame with the translated protein coding sequence (Figure 1B-C). HMA profiles of multiple G0 animals revealed the presence of additional, smaller indels at the 5’ and 3’ CRISPR target regions (Figure 1B, i.e. lanes 3, 8, 10, etc.) as have been observed in other studies.^34,35^ To verify the presence of the smaller indels and the larger in-frame deletions in each animal, we employed a three-step process of HMA, restriction digest, and Sanger sequencing (Figure 1D-E, Supplementary Table 1-2). The majority of smaller indels 10/19 (52.6%) were detected in the 5’ CRISPR target region of G0 animals. Within the 5’ CRISPR target region, there is a unique Hpy81 restriction site that is lost in each of the smaller indels. Consequently, PCR amplification and Hpy81 restriction digestion was used to confirm the presence of the smaller indels in each animal (Figure 1D). Each of these small indels at the 5’ CRISPR target region resulted in premature stop codons in all cases (Figure 1E, Supplementary Table 1). Indels were also observed in the 3’ CRISPR target region in 5/19 (26.3%) animals; however these were not predicted to have an effect on protein translation due to the presence of premature stop codons at the 5’ CRISPR target region.

**Figure 1:**
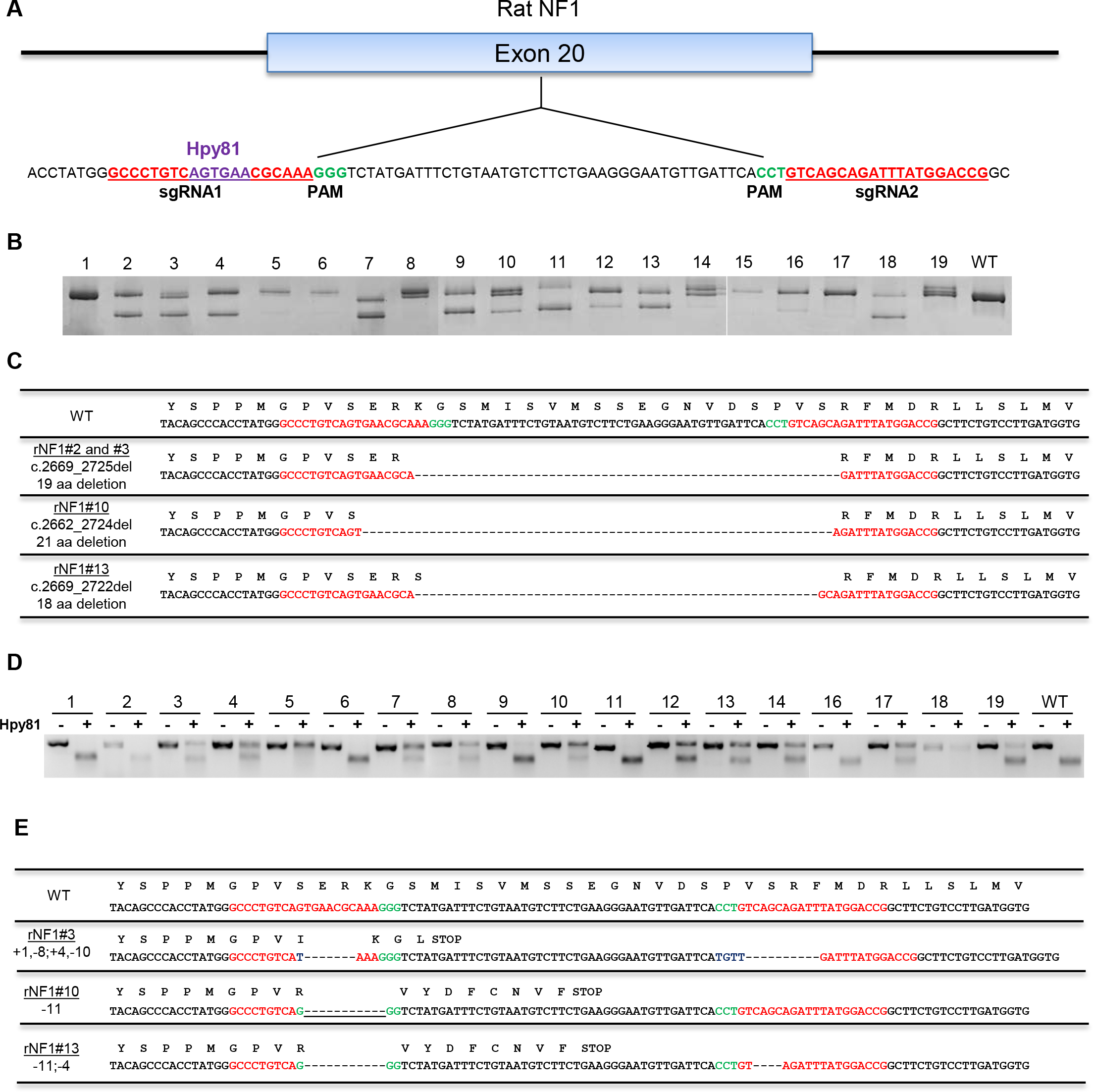
In-frame and premature indels are present in *Nf1* G0 rats. A) CRISPR-Cas9 targeting of the *Nf1* GRD domain in exon 20 using two sgRNAs (red). PAM sequences (green) and the Hpy81 site (purple) used for HMA/restriction digest analysis are shown. B) Heteroduplex PCR amplicons from 19 *Nf1* G0 animals. C) DNA and peptide sequence alignment of the in-frame deletions within *Nf1*exon 20. D) Heteroduplex PCR amplicons from 19 *Nf1* G0 animals were digested with Hpy81 to reveal the presence of smaller indels. No change between the −Hpy81 and +Hpy81 indicates the Hpy81 restriction digestion was lost during CRISPR recombination events. E) DNA and peptide sequence alignment of the small indels at the 5’ and 3’ CRISPR target region.

To investigate the effects of both the *Nf1* in-frame (referred to as **IF**) and premature stop (referred to as **PS**) indels, we bred four founder (F_0_) rats (2 male and 2 female) that carried a mix of indels consisting of in-frame deletions and premature stops with wildtype Sprague-Dawley rats (Figure 1 and Supplementary Table 1). From two male founders we generated two unique lines: ***Nf1***^*IF-57/+*^ from male founder #2 and ***Nf1***^*IF-57/PS-8*^ derived from male #3. From two female founders, the lines ***Nf1***^*IF-63/PS-11*^ (female #10) and ***Nf1***^*IF-54/PS-11*^ (female #13) were generated. Segregation of the IF and PS alleles occurred in the G1 generations for lines ***Nf1***^*IF-57/PS-8*^ and ***Nf1***^*IF-54/PS-11*^ resulting in ***Nf1***^*PS-8/+*^, ***Nf1***^*PS-11/+*^, ***Nf1***^*IF-57/+*^ lines (male #3). These *Nf1* lines were bred out for the following studies.

Upon performing PCR-HMA and sequence analysis, we observed the presence of more than two *Nf1* alleles in individual animals. In ***Nf1***^*IF-63/PS-11*^ we identified 3 *Nf1* alleles: WT allele, an allele with an 11 bp deletion (premature stop), and an allele with a −63 bp deletion (Supplementary Figure 1, Supplementary Table 1). The presence of more than 2 alleles was confirmed independently by HMA and sequencing analysis using unique primer sets in two separate labs. Furthermore, these 3 alleles were transmitted through the germline in G1 and G2 generations and did not segregate in either generation (Supplementary Figure 1). Even though this finding was unanticipated, this is not unique in that the rat genome is known to have additional allelic copies of other genes.^36,37^

### *Nf1* female and male rats develop aggressive mammary adenocarcinomas

*Nf1* mutant rats (including in-frame and premature stop indels) were aged to determine the effects of *Nf1* alterations on tumor development. In *Nf1* female rats, we observed aggressive, multifocal tumorigenesis in the mammary glands at a young age (6-8 weeks). Mammary tumors were observed in 100% (11/11) of G0 females at the average age of 51 days (Supplementary Table 1). Tumor development progressed rapidly in all but one female (*rNf1*#19), whereas the majority of G0 females had to be euthanized before reaching a mature breeding age. We were able to breed two females before their tumor burden required euthanasia (*rNf1*#10 and #13) and support the pups through fostering. Multiple mammary adenocarcinomas (2 - 8) were observed in each animal and were detected in mammary glands 1-5. The detailed tumor pathology for each animal is described in Supplementary Table 3. Notably, two male founders (*rNf1*#2 and #3) developed mammary tumors at 14-16 months of age. Male #2 developed four unique mammary adenocarcinomas, whereas male #3 developed one mammary adenocarcinoma that had to be euthanized due to a larger fibrous histiocytic sarcoma. This aggressive mammary phenotype was highly penetrant and multifocal mammary tumors were observed in the subsequent G1 and G2 generations from each of the *Nf1* lines, including lines derived from male founders (Supplementary Table 3).

Histopathologic analysis of the mammary tumors revealed that both *Nf1* in-frame deletions and premature stop indels induced a wide variety of histopathologic mammary tumor types. Mammary tumors with acinar, solid, ductular, and cystic histology were observed in tumors in *Nf1* female rats from each line (Figure 2A). Moreover, *Nf1* animals often developed mammary adenocarcinomas with mixed histology such as acinar and solid (Figure 2A, left panel); cystic, papillary, ductal, and solid (Figure 2A, middle panel); and cystic, acinar, and solid (Figure 2A, right panel). This diverse mammary histology was observed in G0, G1, and G2 animals. In addition, we observed mixed histopathogy in the mammary adenocarcinomas from male *Nf1* rats (Figure 2B). The level of tumor burden in mammary pads of several animals was substantial, as can be observed in *rNf1* #413 (Supplementary Table 3). As shown in Supplementary Figure 1, one mammary gland from *Nf1* #413 contained 6 separate mammary tumors (2 additional larger tumors were separated for histopathology). Immunostaining of several tumors revealed that the *Nf1* mammary tumors were positive for estrogen receptor, progesterone receptor, and HER2 receptor and highly proliferative based on Ki67 staining (Figure 2C). The results demonstrate the impact of *Nf1* loss of function on mammary tumor initiation in both male and female hormonal environments.

**Figure 2:**
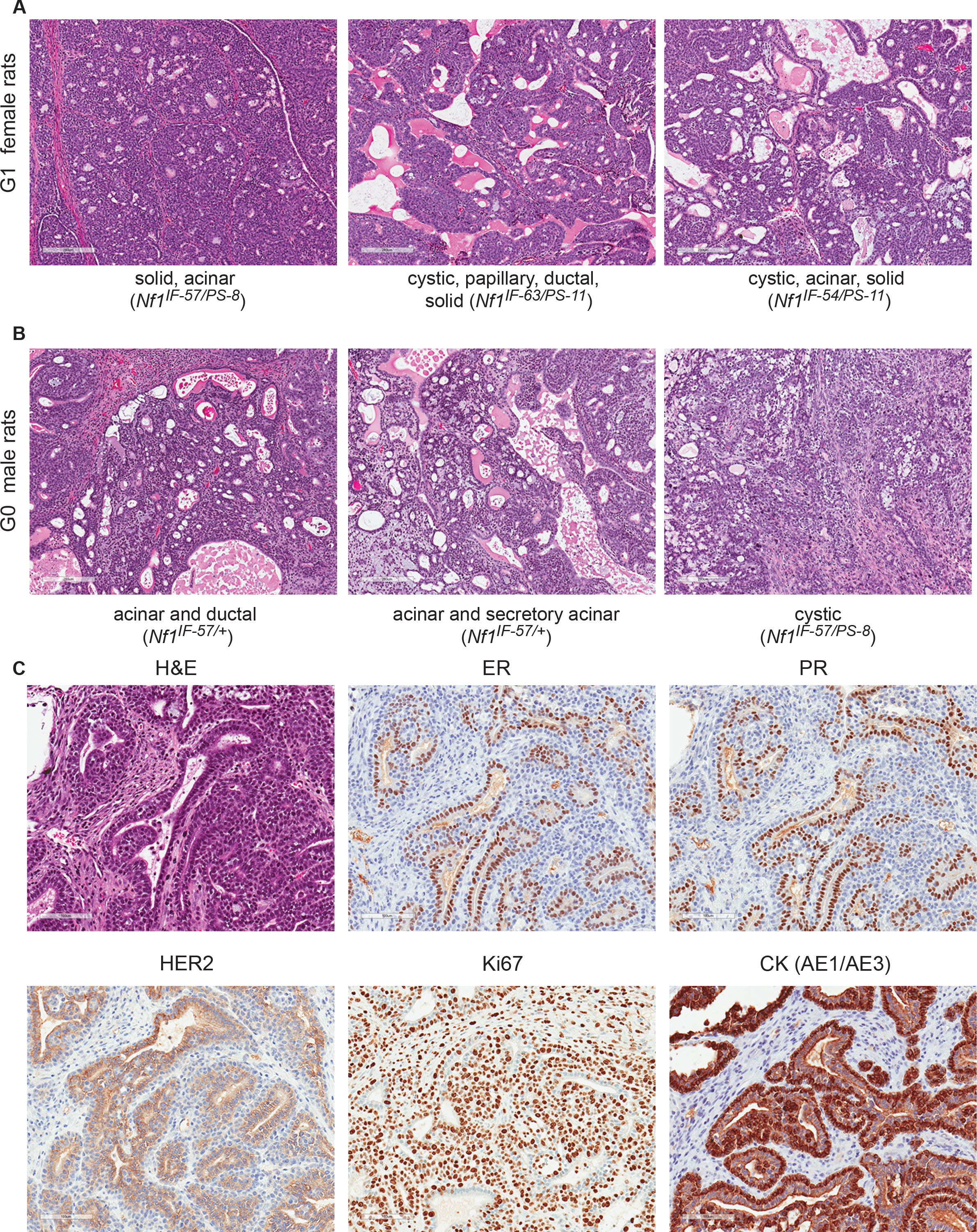
*Nf1* deficiency induces ER^+^/PR^+^ mammary tumors. We observed mammary tumors with diverse mammary histology in F0-G2 animals. A) *Nf1* females and B) *Nf1* males developed mammary adenocarcinomas with mixed histology including acinar, cystic, papillary, ductal, and solid features. All images were taken at 100X magnification. C) Immunostaining of *Nf1* mammary tumors (rNf1 #6) for estrogen receptor, progesterone receptor, HER2, Ki67, and pan-cytokeratin (CK AE1/AE3). All immunostaining images were taken at 200X magnification.

### *Nf1* premature stop indels result in more aggressive tumor progression

To evaluate how in-frame vs. premature *Nf1* indels affect disease burden and survival, we examined the overall survival of G1 heterozygous females. We performed a Kaplan-Meier analysis on overall survival of *Nf1* in-frame (IF) indels (n=35) compared to *Nf1* premature stop (PS) indels (n=24) in three rat lines (*Nf1*^*IF-57/+*^, *Nf1*^*IF-57/PS-8*^, *Nf1*^*IF-54/PS-11*^). Survival analysis of tumor onset demonstrated that there was no statistical difference between the time of onset in premature stop compared to in-frame indels (Figure 3A). Conversely, analysis of disease-specific survival (p<0.06) and overall survival (p< 0.0001) revealed that animals with *Nf1* premature stop indels died due to tumor burden significantly faster than animals harboring *Nf1* in-frame deletions (Figure 3B-C). Notably, all animals in this study died of mammary tumor progression. We compared the overall survival of two *Nf1*^*IF-57*^ lines that were derived from two separate founders (em2 and em3). The divergence in the tumor outcome in the *Nf1* in-frame indels compared to the *Nf1* premature stop indels suggests that the *Nf1* in-frame deletions may result in a partially functional neurofibromin protein and a less aggressive tumor phenotype. Interestingly, we observed that the em2-*Nf1*^*IF-57*^ rats had a significantly shorter survival than em3-*Nf1*^*IF-57*^ rats even though these *Nf1* lines had identical in-frame deletions (p<0.02; Figure 3D). The potential reason for this discrepancy in survival is described below. The fact that tumor initiation was comparable between the *Nf1* in-frame and *Nf1* premature stop indels suggests that a minimal decrease in NF1 function may be sufficient for mammary tumor initiation, yet a more significant loss of NF1 function (as in the *Nf1* premature stop indels) is required for rapid tumor progression.

**Figure 3:**
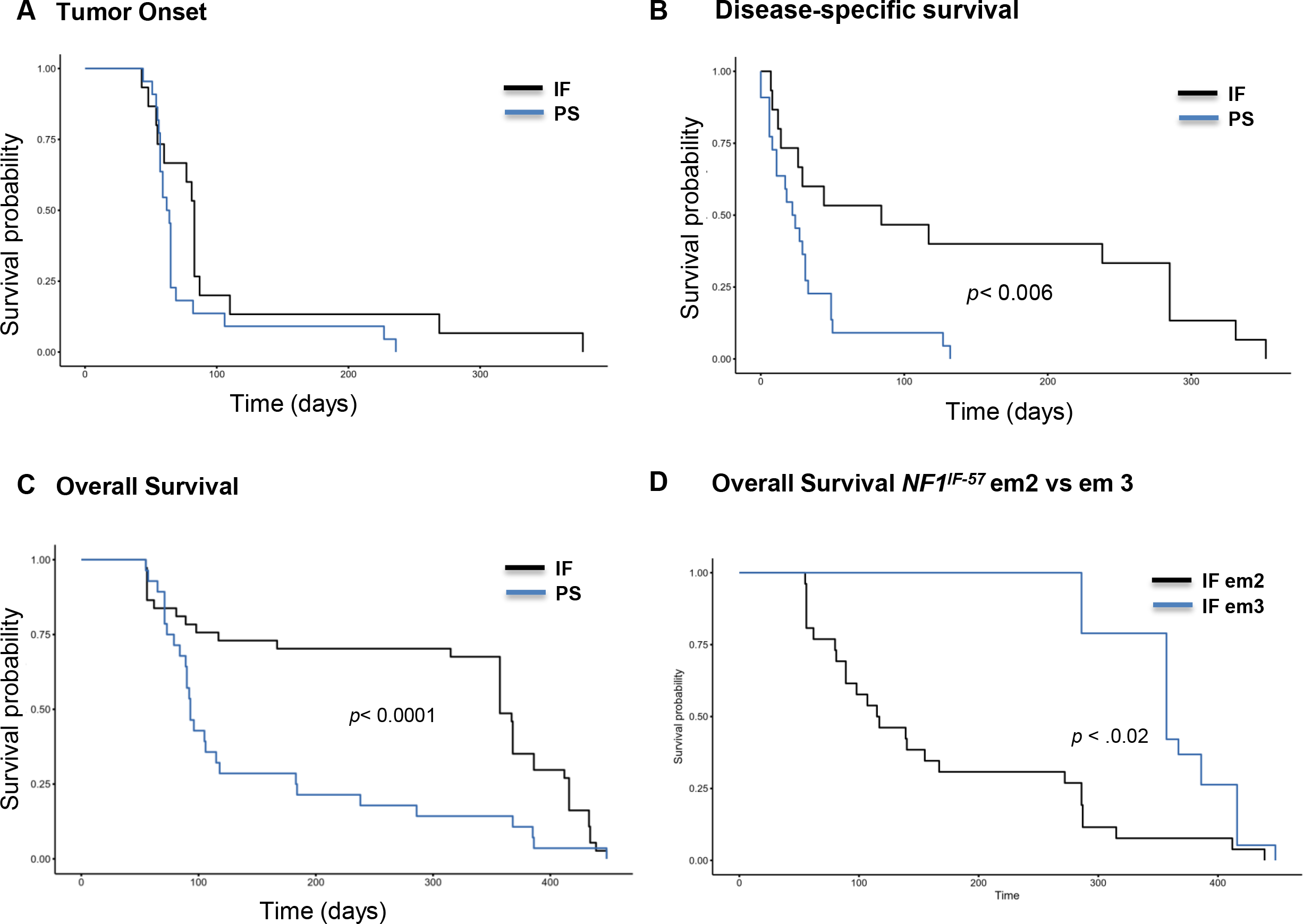
Survival analysis of *Nf1*^IF^ vs *Nf1*^PS^ reveals distinct effects of Nf1 deficiency on tumor onset and survival. Survival analysis was performed on *Nf1*^IF^ (n=35) vs *Nf1*^PS^ (n=24) in the *Nf1*^*IF-57/*+^, *Nf1*^*IF-57/PS-8*^, *Nf1*^*IF-54/PS-11*^ lines. Kaplan Meier plots of A) Tumor onset, B) disease-specific survival, and C) overall survival are shown. D) Overall survival analysis of em2 and em3 lines harboring the *Nf1*^*IF-57/+*^ indel showed a discrepancy in survival. All animals in this study died of mammary tumor progression.

### Distinct neurofibromin isoform is expressed in mammary tissue

To understand how *Nf1* deficiency via mutation of the GRD domain affects neurofibromin expression and activity, we first examined the *Nf1* isoforms that are expressed in distinct *Nf1*^*IF*^ and *Nf1*^*PS*^ lines. Rat embryo fibroblasts (REFs) were isolated from day 13.5 rat embryos and mRNA was isolated. RT-PCR was performed using primers in exon 17 and exons 22-23 (Figure 4A). To differentiate RT-PCR products derived from the WT, IF, and PS alleles we performed restriction digests with Hpy81 endonuclease (Figure 4B). As demonstrated in Figure 1D-E, alleles harboring premature stop indels have lost the Hpy81 restriction site at the 5’ CRISPR site. RT-PCR of WT REFs resulted in a strong band at 826 bp that created two bands at 390 bp and 368 bp with Hpy81 digestion. RT-PCR and Hpy81 digestion of the *Nf1*^*PS-8*^ fibroblasts resulted in *Nf1*^*WT*^ bands and an undigested band at 745 bp. Expression of the WT allele was observed in each of the *Nf1* in-frame and premature stop REF lines. We compared the em2-*Nf1*^*IF-5*^ and em3-*Nf1*^*IF-57*^ lines that had distinct survival curves (Figure 3D). Even though genotyping demonstrated both of these lines have a 57 bp deletion in exon 20, we observed distinct *Nf1* mRNA isoform expression in these lines (Figure 4B, notated with *). To determine the difference in the *Nf1*^*IF-57*^ mRNA isoforms, we cloned and sequenced each of the products. Overall, we sequenced seventeen clones where the majority contained the predicted *Nf1*^*IF-57*^ and *Nf1*^*WT*^ isoforms, but we identified 3 unique clones harboring exon 21 deletions or extended exon 20 deletions resulting in *Nf1* premature stops. For example, one mRNA isoform contained the −57 bp in-frame deletion in exon 20 and an additional 140 bp deletion in exon 21 (Figure 4C). This exon 21 deletion mRNA was exclusive to the em2-*Nf1*^*IF-57*^ line and resulted in a premature stop. These results substantiate the requirement of a functional GRD domain within NF1 to suppress tumor progression.

**Figure 4:**
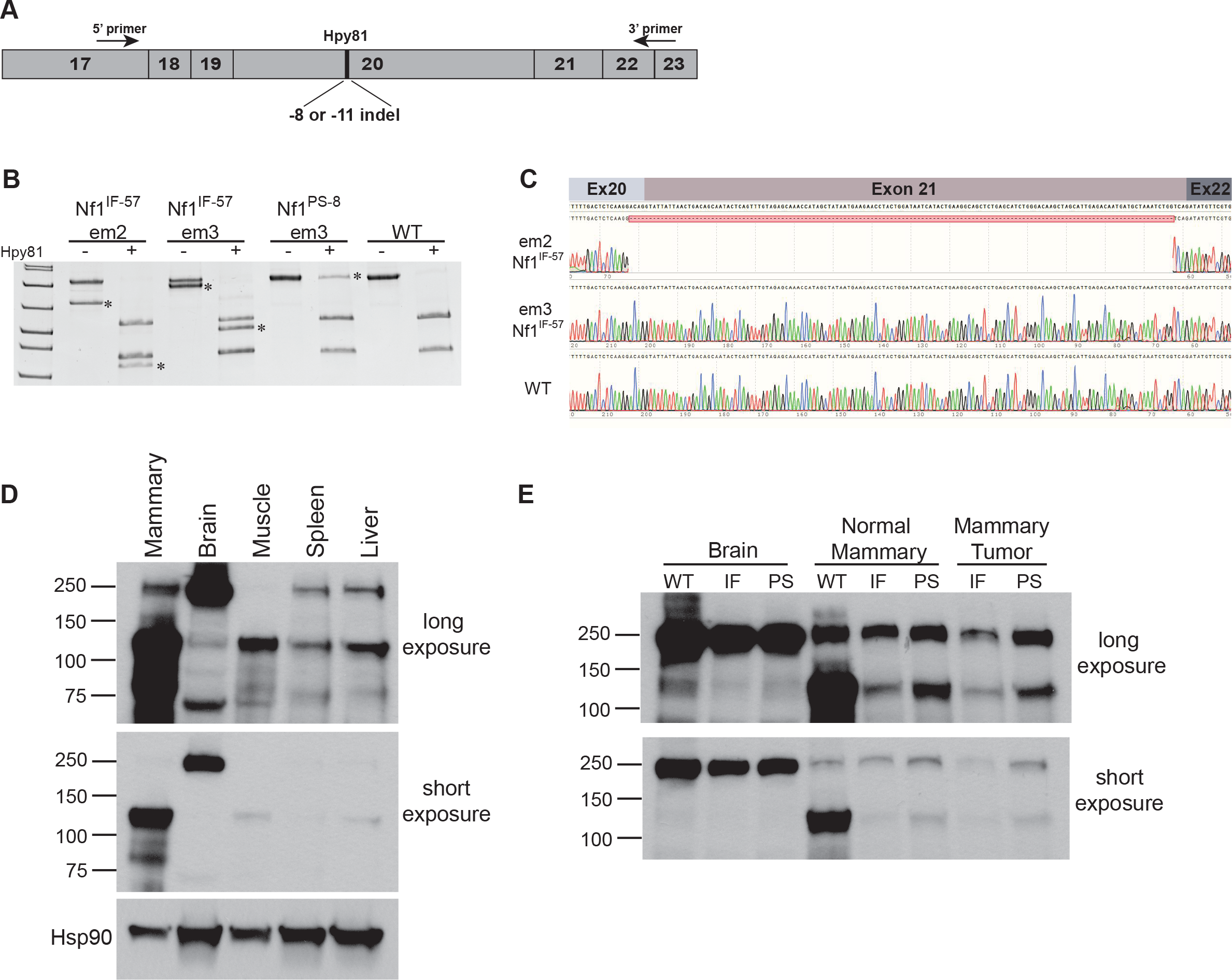
Analysis of *Nf1*^*IF*^ and *Nf1*^*PS*^ mRNA and protein isoforms. A) Schematic of RT-PCR analysis of *Nf1* exons 17-23. B) RT-PCR products were digested with Hpy81 and separated by PAGE to differentiate *Nf1*^*IF*^ and *Nf1*^*PS*^ mRNA. Unique bands were observed in the em2 and em3 *Nf1*^*IF-57*^ REF cDNA (noted by *). C) Sequence analysis of em2-*Nf1*^*IF-57*^ and em3-*Nf1*^*IF-57*^ REF cDNA. Immunoblot analysis showing D) tissue-specific expression of rat neurofibromin isoforms and E) differential expression of the neurofibromin 250 and 125 kDa bands in matched brain, normal mammary tissue, and mammary tumor at both a long (top) and short (bottom) exposure.

Several unique isoforms have been characterized in the *Nf1* gene with the main isoform identified at 250 kD; however the majority of these studies have been performed in brain, neuronal, and muscle tissues.^38–40^ To our knowledge, neurofibromin expression has not been previously evaluated in the mammary glands of humans or rodents. To understand the effect of *Nf1* deficiency in the mammary gland, we first performed Western blot analysis of neurofibromin in the normal mammary, brain, muscle, spleen, and liver of adult rats and observed distinct isoform expression patterns among the tissue types (Figure 4D). Even though the established 250 kD neurofibromin isoform is highly expressed in brain tissue, its expression was significantly less in the other tissues including the mammary gland. In the mammary gland, the predominant neurofibromin isoform was observed at 125 kD. This 125 kD isoform was also observed in the muscle, spleen, and liver, yet minimally expressed in the brain. In Figure 4D, we compared neurofibromin expression in the brain, normal mammary glands, and mammary adenocarcinomas from *Nf1*^*WT*^, *Nf1*^*IF*^, and *Nf1*^*PS*^ rats. In the brain, the 250 kD isoform was highly and equally expressed in the *Nf1*^*WT*^, *Nf1*^*IF*^, and *Nf1*^*PS*^ animals. In the normal mammary gland there was no difference in expression of the 250 kD isoform between the *Nf*^*WT*^, *Nf1*^*IF*^, and *Nf1*^*PS*^ animals, however there was a substantial decrease in expression of the 125 kD isoform in both the *Nf1*^*IF*^ and *Nf1*^*PS*^ mammary glands compared to WT mammary glands. In mammary adenocarcinomas, 125 kD isoform was further decreased in both the *Nf1*^*IF*^ and *Nf1*^*PS*^ tumors and there appeared to be a slight loss of the 250 kD isoform in the *Nf1*^*IF*^ mammary tumor. Overall these results demonstrate the distinct neurofibromin isoform tissue preferences that are present and the inverse relationship of the 250 kD vs. 125 kD isoforms in the brain and mammary tissues.

### *NF1* is commonly deleted in human sporadic breast cancer and correlates with ER networks

Even though it is known that neurofibromatosis patients have a significantly increased breast cancer risk and that *NF1* is mutated in sporadic breast cancers, the impact of NF1 deficiency and RAS deregulation in sporadic breast cancer is often overlooked. To interrogate the impact of *NF1* loss in breast cancer we performed an analysis of the METABRIC breast cancer dataset which contained 2051 patients with clinical annotation including CNV and SNP genotypes.^33^ In this cohort, there were 43 patients (2.1%) with truncating mutations, including frameshift deletions or insertions, nonsense mutations, and splice site alterations (Figure 5A). Notably, these mutations occurred throughout the *NF1* gene similar to the mutation diversity observed in neurofibromatosis patients. After removing patients with *NF1* SNP mutations, we analyzed *NF1* copy number alterations (CNAs) and identified CNAs in 32.9% of patients, with 24.5% having *NF1* shallow deletions (defined as potential heterozygous deletions). To determine the effect of *NF1* shallow deletions on survival, we conducted a survival analysis using the Cox proportional hazards model with random effects (frailty model). When accounting for ER status and age at diagnosis, a patient with a *NF1* shallow deletion is 1.65× (p < 1.6e-05) more likely to die within the first 10 years compared to patients with diploid *NF1* status (Figure 5B).

**Figure 5:**
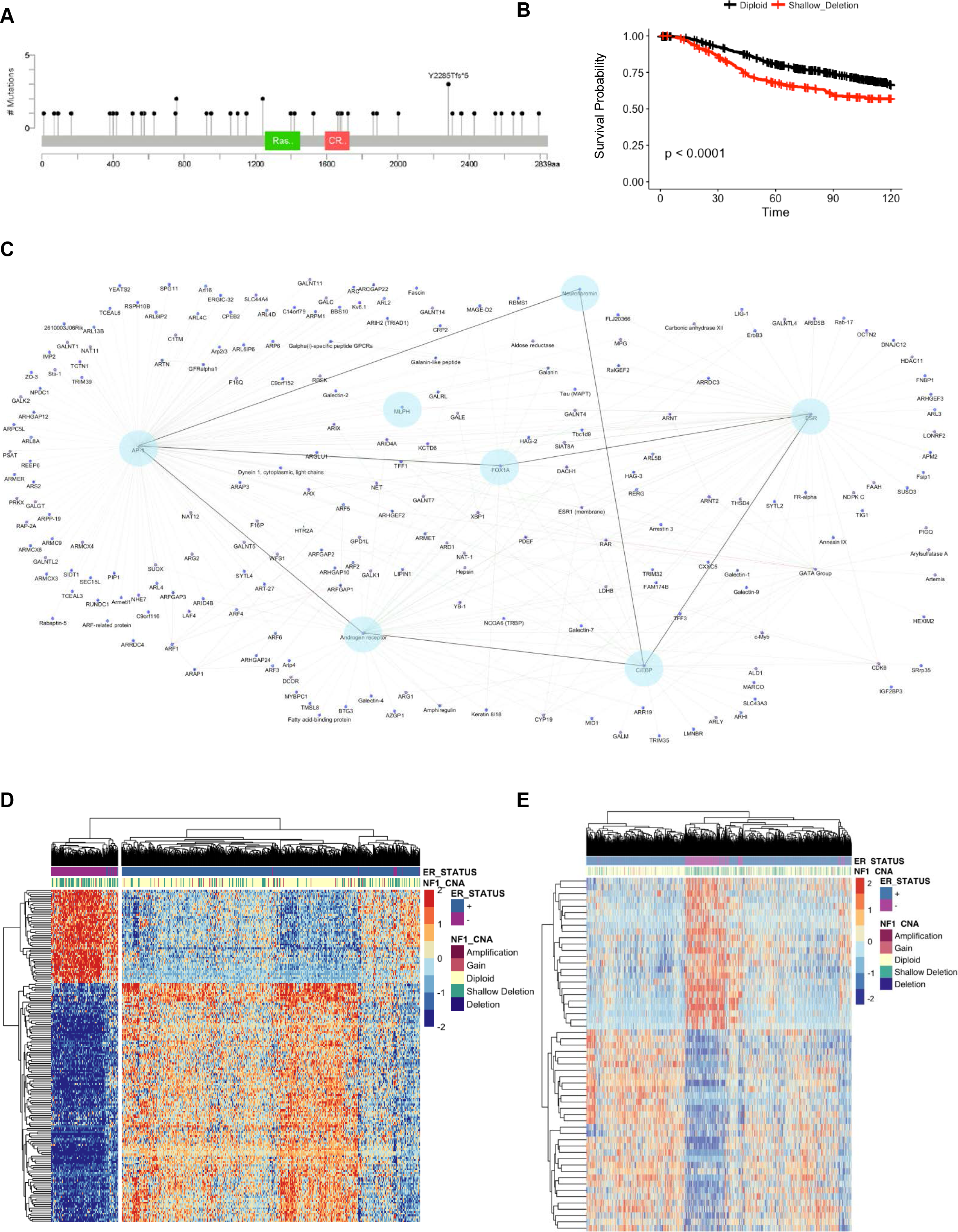
*NF1* shallow deletions are frequently present in sporadic breast cancers and associate with ER and FOX1A. A) Truncating mutations were detected throughout *NF1* in METABRIC patients (cBioportal) B) Survival analysis of breast cancer patient survival comparing *NF1* diploid and shallow deletion copy number status. C) Network visualization of WGCNA module genes associated with *NF1* shallow deletion copy number status. D) Heatmap of METABRIC gene expression data of WGCNA module genes correlated with *NF1* copy number and ER status. D) Heatmap of METABRIC gene expression data of WGCNA module genes associated with RAS activation signatures.

To identify gene expression networks that correlate with *NF1* copy number status, we utilized weighted gene co-expression network analysis (WGCNA).^41^ From the METABRIC dataset, expression data were analyzed for patients with available copy number and gene expression data (n=1427). Since *NF1* is in close proximity to *HER2* on chromosome 17 and is often co-amplified, we removed HER2-amplified patients from this analysis. From the 18,049 genes in the METABRIC dataset, 2,218 were placed in 12 co-expression modules. Modules contained between 30-500 genes, with an average module size of 200 genes. Interestingly, one module (179 genes) associated with *NF1* shallow deletion copy number status contained several genes that are considerably important in both ER+ breast cancers and endocrine resistance, including *ESR1* (Figure 5C). Hub genes for this module, or genes with high connectivity, included *FOXA1* and *MLPH*. *FOXA1*, a Forkhead family transcription factor, regulates ER binding and transcriptional activity.^42^ *FOXA1* expression correlates with luminal subtype A breast cancer and is a significant predictor of cancer-specific survival in ER+ tumors.^43^ *MLPH* expression is also associated with longer survival in breast cancer.^44^ Further network visualization of genes within this ER-associated co-expression network revealed additional connections with *AP1*, *AR* (androgen receptor) and *C/EBPα*. AP1 is also known as the transcription factor JUN. RAS/ERK signaling is known to regulate JUN at the transcriptional and post-translational (phosphorylation) level and can modify lysine acetylation in JUN DNA binding regions.^45^ Recent studies have demonstrated that *AR* is expressed in 77% of breast cancers (88% ER+, 59% HER2+, 32% TNBC)^46^ and is involved in endocrine resistance in ER+ breast cancers.^47,48^ Although there are studies demonstrating interaction of C/EBP (CCAAT/enhancer binding protein) at ER transcriptional binding sites, the consequence of C/EBP activity in breast cancer is unclear.

Unsupervised clustering of genes from this WGCNA module with METABRIC clinical data indicates a strong association with patient ER status, specifically ER-negative breast cancer (Figure 5D). To understand if shallow deletions of *NF1* correlate with deregulated RAS signaling, we employed expression signatures developed from KRAS-mutant cancers and performed a secondary WGCNA analysis and unsupervised clustering with 788 genes related to ER, RAS and KRAS signaling pathways. This analysis distinguished three modules associated with *NF1* copy number status, with one module associated specifically with *NF1* shallow deletions (Figure 5E). These results validate the presence of RAS activation in *NF1*-related breast cancers. Moreover, our WGCNA analyses revealed novel connections between NF1 deficiency, RAS signaling, and ER signaling.

### *Nf1*-deficient tumors are estrogen-dependent

To examine whether our *Nf1* deficient rat breast cancer model was estrogen dependent, we performed ovariectomies on rats containing at least one tumor greater than 1500 mm^3^. Ovariectomies were performed on 5 rats with multiple tumors from three distinct Nf1 lines (Figure 6F and Supplementary Figure 3). Upon ovary removal, tumor size diminished at a surprisingly rapid pace (mean reduction per day = 5.33%; CI [4.89, 5.78]; p < 0.000000001). Tumor reduction was observed in all of the tumors regardless of Nf1 status (in-frame vs. premature stop indels) and the mean total percent reduction in tumor volume was 90.95%; (CI [78.79, 97.28], p = 0.0000018) (Figure 6G and Supplementary Figure 3). These results confirm estrogen-dependency in the rat *Nf1*-deficient breast cancer model and support the biological significance of the NF1-ER networks that were identified by WGCNA.

**Figure 6:**
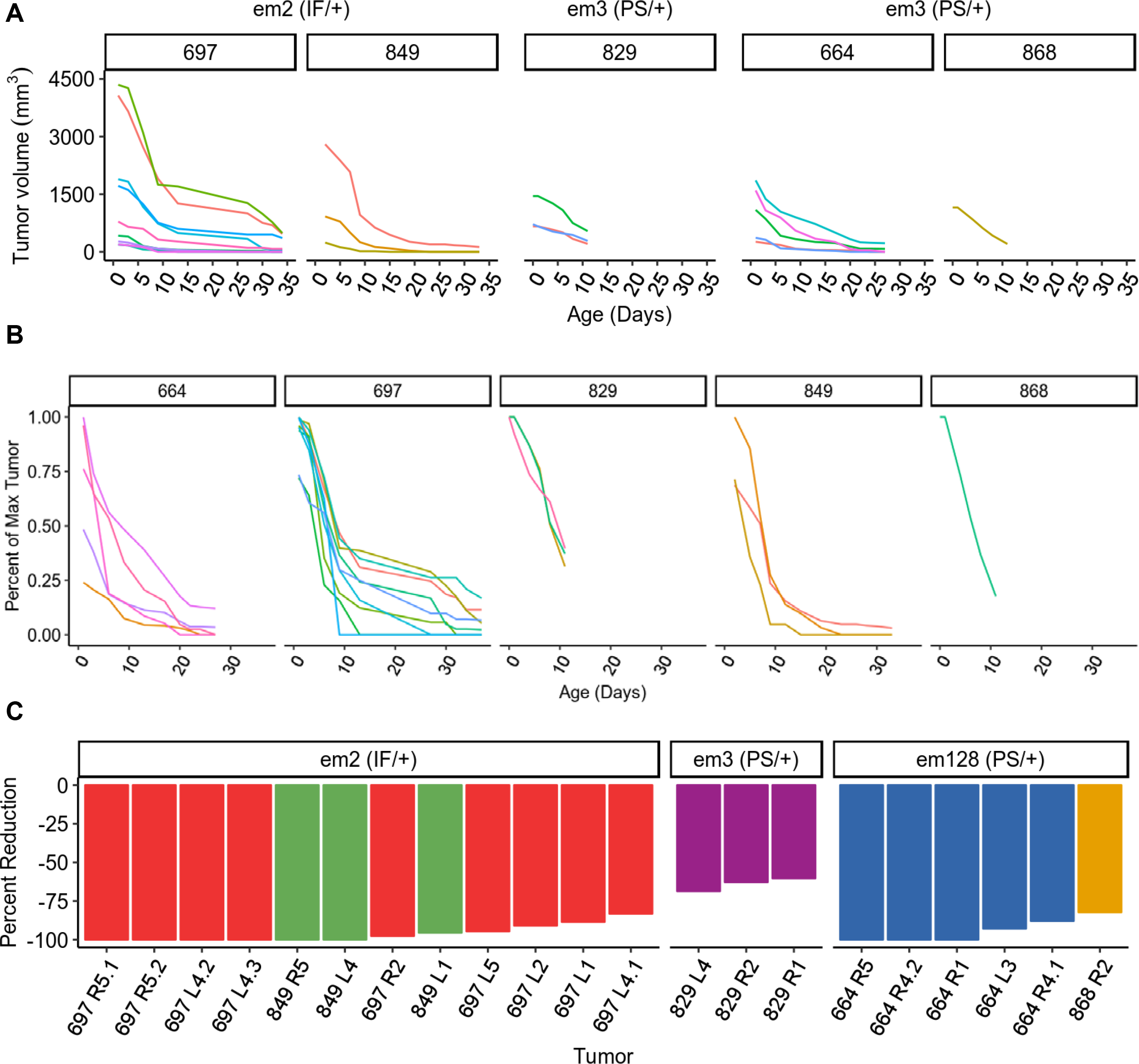
Estrogen depletion via ovariectomy results in rapid tumor regression in *Nf1*-deficient tumors. A) Tumor volume was plotted after ovariectomies (Day 0) for individual *Nf1*^*IF*^ and *Nf1*^*PS*^ rats. B) Percent tumor change over time of multiple tumors in individual *Nf1*^IF^ and *Nf1*^PS^ rats after ovariectomy (Day 0). C) The percent reduction in tumor size for all tumors from 3 Nf1 rat lines is shown. Distinct colors are used to represent individual rats and the mammary pad location of each tumor is notated. Note that several rats had multiple measurable tumors within an individual mammary pad.

## Discussion

The RAS signaling pathway is intricately involved in the initiation and progression of numerous cancer types; however the mechanism by which RAS is deregulated in breast cancer is unclear. Here we present data that illuminates the role that *NF1* plays in breast cancer and a novel model for interrogating the interaction between NF1 and ER in both sporadic and NF1-related breast cancer. Even though NF patients develop a variety of cancers, the high breast cancer risk in women with neurofibromatosis was not been highlighted until recently.^15–17^ *NF1* was firsts implicated in sporadic breast cancer by the initial comprehensive genome breast cancer studies where *NF1* mutations were identified in 2-4% of breast cancers.^18,19^ Our analysis of the extensive METABRIC dataset verified the *NF1* mutation frequency, but also discovered *NF1* shallow deletions in 25% of sporadic breast cancers. This verifies the *NF1* shallow deletions that have been identified in the TCGA breast cancer dataset.^20^ Most importantly, our analysis revealed that *NF1* shallow deletions correlated with poor outcome after the first 10 years. One clinical challenge for oncologists is determining which patients primary breast cancer should be treated aggressively. Our results indicate that *NF1* copy number may be a potential prognostic indicator in breast cancer. In addition to *NF1*, the RasGAP gene *RASAL2* is altered in a sporadic breast cancer. McLaughlin et al. demonstrated that *RASAL2* expression is substantially decreased in luminal B breast cancers and low *RASAL2* expression correlates with metastasis and poor survival.^49^ Our WGCNA analysis revealed an unknown network between NF1, ER, and RAS signaling in breast cancer. Together, these studies indicate that loss of RAS suppression may be more important in breast cancer than activation of RAS signaling through mutation or amplification.

Together our CRISPR rat *Nf1* model and our WGCNA analysis of human breast cancer indicate that NF1 is intricately connected to ER networks, including several genes that have been directly linked to endocrine resistance (i.e FOXA1 and AR). The rate of tumor regression in the *Nf1* tumors after estrogen ablation was remarkably rapid and the fact that all of the *rNf1* tumors diminished regardless of mutation (PS or IF) suggests that *Nf1*-deficienct tumors are universally estrogen-dependent. Interrogating the mechanisms by which NF1 interacts with ER will involve a careful analysis both transcriptional and signaling interactions. NF1 is a large protein and there is a limited knowledge of NF1 protein-protein interactions besides the RAS interactions at the GRD domain. It is possible that NF1 may directly interact with genes involved identified in the NF1-ER network at a transcriptional level, yet indirect signaling NF1-RAS-ER signaling interactions are also likely since RAS signaling regulates JUN (AP1) at the transcriptional and post-translational level.^45^ A recent study that assessed driver genes during the genomic evolution of breast cancer metastasis revealed that *NF1* mutations substantially increase in recurrent breast cancers, but this increase was only observed in ER-positive breast cancers^50^. This findings in addition to 1) the correlation of *NF1* shallow deletions with poor outcome, 2) network connectivity between NF1-ER-FOXA1-AR, and 3) estrogen-dependency in the Nf1 tumors, indicate that NF1 may be a key player in endocrine reistance.

The *Nf1* rat model also revealed some unexpected alternative mRNA and proteins isoforms that underscore our limited understanding of NF1 expression. For example, we observed two distinct neurofibromin isoforms that are differentially expressed in the brain and mammary gland. Both in-frame and premature stop sequence variants resulted in significantly reduced neurofibromin mammary expression, whereas expression of the 250kD isoform was preserved in the brain regardless of *Nf1* status. It is unclear why the wild type allele was unable to effectively compensate for heterozygous loss of NF1 function in the mammary gland. Collectively, these findings suggest that RAS regulation in the breast is distinct from other tissues, and that breast cancer predisposition may be linked to a unique isoform of neurofibromin that is abundantly expressed in mammary tissue.

One issue that vexed us initially was the divergence of survival between two genotype-matched *Nf1*^*IF-57*^ lines. Analysis of the mRNA in em2 *Nf1*^*IF-57*^ and em3 *Nf1*^*IF-57*^ lines identified an additional deletion in exon 21 in em2 *Nf1*^*IF-57*^. Differential mRNA isoform expression has been observed in breast cancer and may occur through alternate promoter usage, alternate splicing, and alternate 3’UTR usage.^51^ Importantly, differential mRNA isoform expression has been identified and associated with distinct breast cancer subtypes. In NF patients, there is often a discrepancy between genotype and phenotype. For example, one NF patient may develop numerous cutaneous and plexiform neurofibromas, where another patient with an identical *NF1* mutation may have few cutaneous neurofibromas and develop breast cancer at a young age. Our data raises the possibility that *NF1* isoform expression may play a role in these clinical inconsistencies.

In summary, we have developed a novel *Nf1* rat model is invaluable for interrogating deregulated RAS signaling and ER networks in sporadic and NF-related breast cancer. Moreover, we identified a frequent mechanism for RAS deregulation through *Nf1* shallow deletion that is present in 25% of sporadic breast cancers. The correlation of *Nf1* shallow deletions with poor prognosis and ER-FOXA1-AR networks indicates that NF1 may be an important prognostic indicator and therapeutic target in endocrine-resistant breast cancers.

## Methods

### CRISPR/sgRNA design and synthesis, and Cas9 mRNA synthesis

CRISPR targets were identified in exon 20 of the rat *Nf1* gene (NCBI rn5 rat genome) using an online design tool (crispr.mit.edu). A region near the middle of exon 20 was targeted by one guide sequence (5’ CRISPR), GCCCTGTCAGTGAACGCAAA (*GGG*), while a downstream region was targeted by a second guide sequence (3’ CRISPR), CGGTCCATAAATCTGCTGAC (*AGG*). The rn5 rat genome reference sequence and annotation for CRISPR target design.

### Rat breeding, microinjection into zygotes and blastocyst culture

Sprague-Dawley rats were purchased from Charles River Laboratories. Female rats were superovulated with injection of 20 IU PMSG, followed by injection of 50 IU HCG and immediate mating to CD Sprague Dawley studs 48 hours later. The pronuclei of fertilized rat zygotes were injected with 40 ng/μL Cas9 mRNA and 20 ng/μL each of two sgRNAs. Surviving eggs were surgically transferred to CD Sprague-Dawley pseudopregnant females. All animal experiments were performed in accordance with institutional IACUC protocols.

### Detecting the presence of indels by PCR-HMA and amplicon sequencing

Genomic DNA from rat-tail biopsies was obtained by phenol extractions. DNA samples were directly used as a PCR template to amplify a region flanking the CRISPR target site. PCRs were set up using the following oligonucleotide primers: Ex20-F 5’-TCA ACA TGA CTG GCT TCC TC–3’ and Ex20-R 5’-CAT TGG ATA CAG AGC AGG ACT C–3’ to obtain a 284 bp fragment. The amplicons were subjected to denaturation-slow renaturation to facilitate formation of heteroduplexes using a thermocycler. These samples were then resolved on polyacrylamide gels (6%) and the resulting mobility profiles used to infer efficiency of CRISPR-Cas9 nuclease activity. PCR using same primers flanking CRISPR target site (Ex20-F and Ex20-R) was performed to amplify regions with indels (284 bp). The amplified products were cloned using the TOPO-TA cloning kit (Invitrogen, Carlsbad, CA). Ten representative colonies were picked from each plate and grown in 4mL liquid cultures to isolate plasmid DNA. Plasmid DNA was sequenced using M13F and M13R primers.

### Genotyping

Rat DNA was collected from tail clips by phenol extraction (Invitrogen 15593-031). Target regions of the DNA were PCR amplified using the forward primer rNF1-1A 5’– CTT AGG CTG CAG AAA GTC TTC – 3’ and reverse primer rNF1-1B 5’– CTT CAC CTG TCC TTG AGA GTC – 3’. PCR products were digested with Hpy8I (Thermo Fisher Scientific). After digestion, DNA fragments were separated by electrophoresis on a 2% agarose gel and stained with ethidium bromide. Premature stop (439 bp) and in-frame (185 bp) mutations were identified by gel imaging.

### RT-PCR

RNA was isolated from rat tissue by Trizol extraction (Invitrogen). RNA was treated with DNase I (Thermo Fisher Scientific #18068015). Complementary DNA (cDNA) was preapred by using random primers (Life Technologies 48190-011) and SuperScript™ II Reverse Transcriptase (Invitrogen 18064-022). PCR was performed with forward primer (5’ – TCAGTACACAACTTCTTGCC – 3’) and reverse primer (5’ – GAACACGAACATATCTGACC – 3’) to create a 826 bp amplicon. These PCR products were digested with the restriction enzyme Hpy8I as described above to identify mutations. Digested PS amplicons make a 745 bp product and digested wildtype make a 390 bp and 368 bp product. Digested PCR products were ran on a 10% TGX-PAGE gel (BioRad) and stained with ethidium bromide.

### Histology

Tumors were fixed and processed following the previous methods ^52^. DAKO PT Link was used for antigen retrieval. Immunohistochemistry staining was done with ER (Thermo MA5-13304), HER2 (Thermo MA5-13105), Cytokeratin (DAKO AE1/AE3), and PR (Thermo MA1-411) using the DAKO Autostainer Link 48. Ki67 (Spring Bioscience SP6) was stained using the Ventanna Discovery Ultra.

### Rat Embryo Fibroblast (REF) Isolation

At embryonic day 13.5 embryos were harvested and isolated independently in petri dishes with PBS. The placenta was removed followed by the yolk sac. Embryos were then dissected by removing the head, organs, and blood vessels. The remaining tissue was transferred to a new petri dish and finely chopped using surgical blades. Tissue was then digested in 0.25% trypsin for 15 minutes at 37C. Cells were passed through a cell strainer to isolate single cells. Cells were plated and grown in DMEM (Gibco 11995-065) supplemented with 10% FBS and 1% penicillin.

### Immunoblot Analysis

Rat tissue was homogenized with a hand held pellet pestle in RIPA buffer (20 mM TrisHCL, pH 7.6; 5 mM EDTA; 150 mM NaCl; 0.5% NP-40; 50 mM NaF; 1 mM beta-glycerpphosphate) supplemented with PhosSTOP (Roche) and protease inhibitors (Roche). Samples were resolved by SDS-polyacrylamide gel electrophoresis on a 7.5% TGX-PAGE gel (BioRad) and transferred overnight at 4°C. Immunoblotting was performed using the following antibodies: neurofibromin H-12 (SC-376886) and D (SC-67) from Santa Cruz Technology; McNFn27b #MA1-085 from Thermo Scientific; β-actin (#3700), HSP90 (#4877) and GAPDH (#2118) from Cell Signaling Technology.

### Statistical Analysis

For analysis of *Nf1* rat survival, data were plotted using Kaplan-Meier curves and p-values were calculated using a Cox mixed-effects model with a frailty term for litter (https://CRAN.R-project.org/package=coxme). Linear contrasts with Benjamini-Hochberg false discovery rate adjustments were used to test specific hypotheses. All analyses described below were performed using R v 3.4.0. (https://cran.r-project.org/) All hypotheses were two-sided with a significance level of 0.05. Ovariectomy analysis was analyzed using a multi-level mixed-effects beta regression with a random intercept for tumor, clustered within rat lines. This model was fit using the R package glmmTMB^53^ and the squeeze algorithm.^54^

### METABRIC analysis

Survival analysis was conducted in R v(3.4.2) using the packages survival (https://cran.r-project.org/web/packages/survival/index.html), survminer (https://cran.r-project.org/web/packages/survminer/index.html), and coxme. Clinical, copy number and gene expression data from the METABRIC dataset^33^ were accessed using the cgdsr (https://cran.r-project.org/web/packages/cgdsr/index.html) package in R. Patients were selected based on the availability of both copy number and gene expression data, and HER2 negative status. Gene-co expression network analysis was conducted using the WGCNA (v1.61) package in R to identify gene coexpression modules from mRNA expression data (Illumina Human v3 Microarray).^41^ Data were normalized as previously described. Unsigned correlations were used with a soft threshold value β of 10 and a treecut value of 0.15, a minimum number of genes in the module set to 30. The β and treecut parameters were chosen after assessing the quality of modules detected, and all other parameters used default settings. Unsupervised clustering of WGCNA results was run using the pheatmap package in R (v1.0.8). Specific gene networks were visualized using MetaCore from Thomson Reuters (v6.33 build 69110). The RAS and KRAS gene lists used in the analysis were provided by the Broad Institute MSigDB database (v6.1) Hallmark gene set collection and BioCarta (c) (2000-2017 BioCarta, all rights reserved).

## Acknowledgements

We dedicate this work in honor of Patricia Graveel and all neurofibromatosis patients dealing with breast cancer. The authors would like to thank Dr. Ben Johnson and Jamie Grit for their critical review of this manuscript, Dr. Casey Droscha and Dr. Bart Williams for assistance with CRISPR designs, and also Lisa Turner (VAI Histology) and the VAI Vivarium staff for their strong work on this project. Funding for this research was made possible by the Breast Cancer Research Foundation (BCRF-17-159) and the VARI Faculty Innovation Award.

## Author Contributions

Study conception and design: PSD, CRG, MRS; acquisition and analysis of data: PSD, EAT, EJE, ZM, EEG, MC, ANT, AKC, TK, RTB, MJB, CRG, MRS; provided reagents: BE, RAK; drafting of manuscript: CRG, MRS; critical revision: PSD, EAT, ZM, MJB; CRG and MRS are the guarantors of this study.

## ADDITIONAL INFORMATION

**Supplementary information** is available at the NPJ Breast Cancer website.

**Competing interests:** The author declares that they have no competing financial interests.

---

**Supplementary Figure 1:**
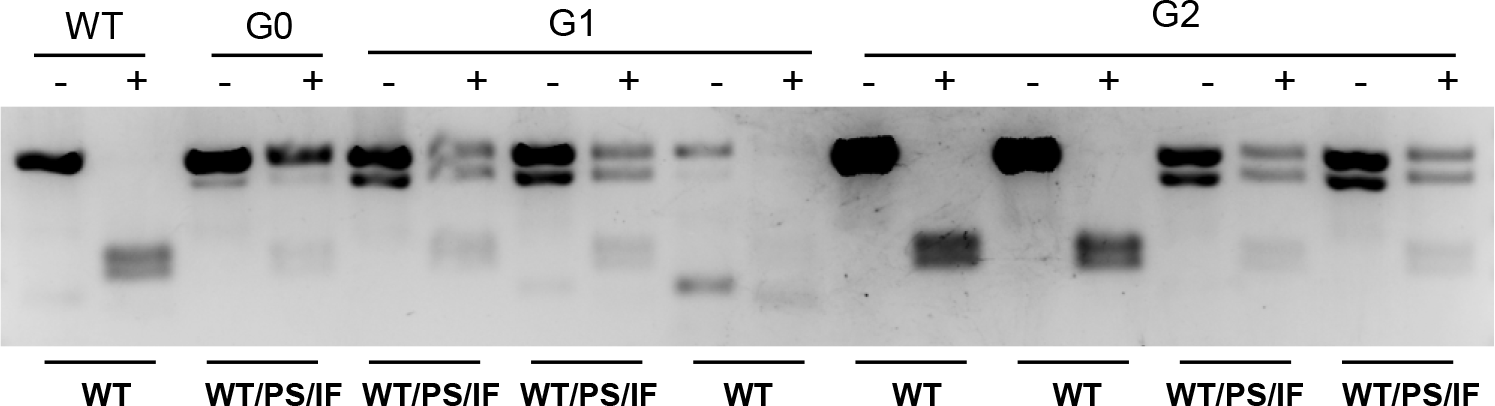
Three Nf1 alleles identified in *Nf1*^*IF-63/PS-11*^ line. HMA and sequencing identified 3 *NF1* alleles in *Nf1*^*IF-63/PS-11*^ : a WT allele, an allele with an 11 bp deletion (premature stop), and an allele with a -63 bp deletion. The presence of more than two alleles was confirmed using unique primer sets in two separate labs. The three alleles were transmitted through the *Nf1*^*IF-63/PS-11*^ germline in F1 and F2 generations and did not segregate in either generation as shown in the HMA profiles of individual F1 and F2 *Nf1*^*IF-63/PS-11*^ rats.

**Supplementary Figure 2:**
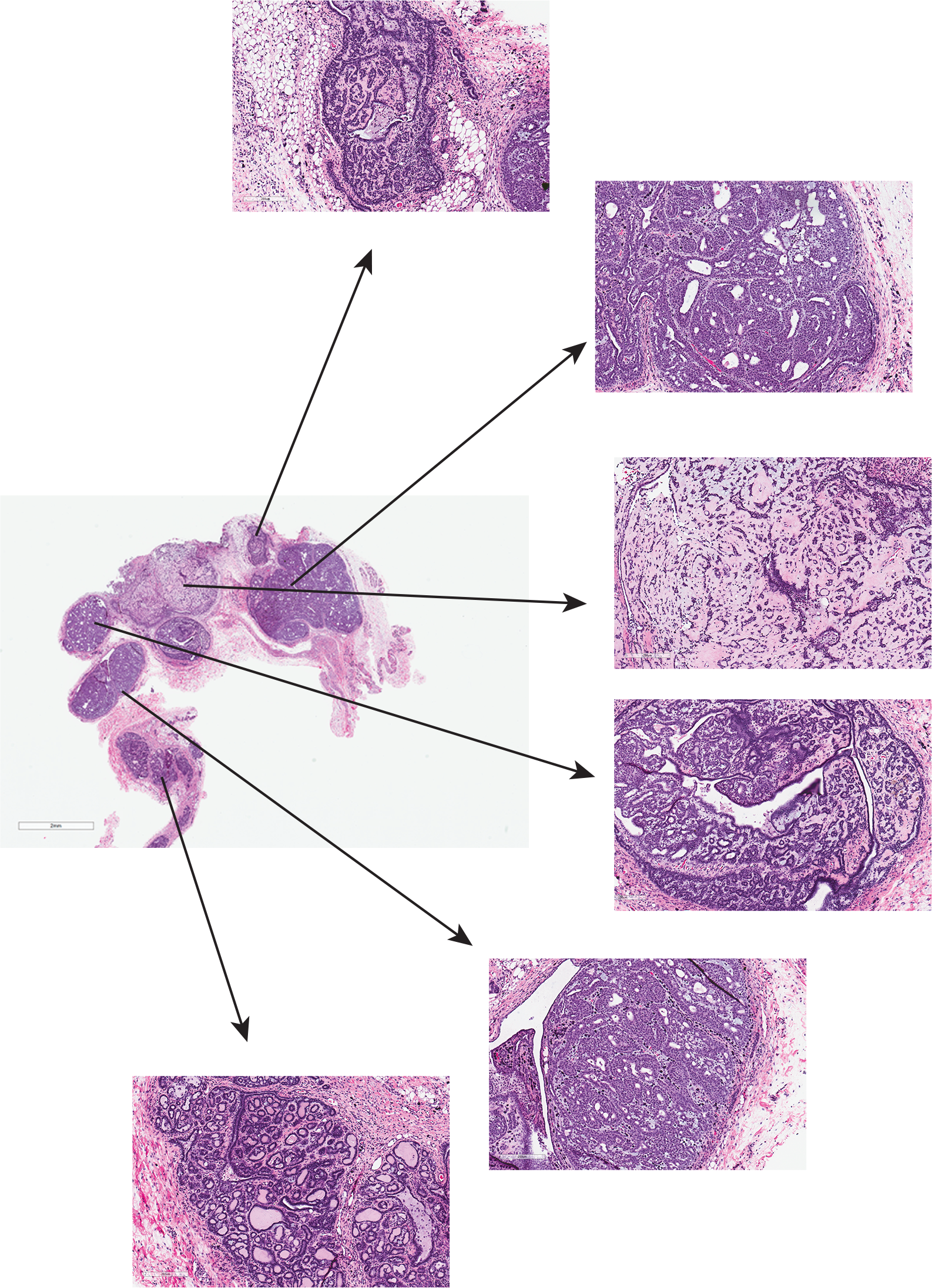
Multiple tumors are commonly observed in *Nf1* mammary glands. The level of tumor burden in the mammary pads of several animals was substantial. Shown above is the 4th mammary pad from *rNf1* #413.

**Supplementary Figure 3:**
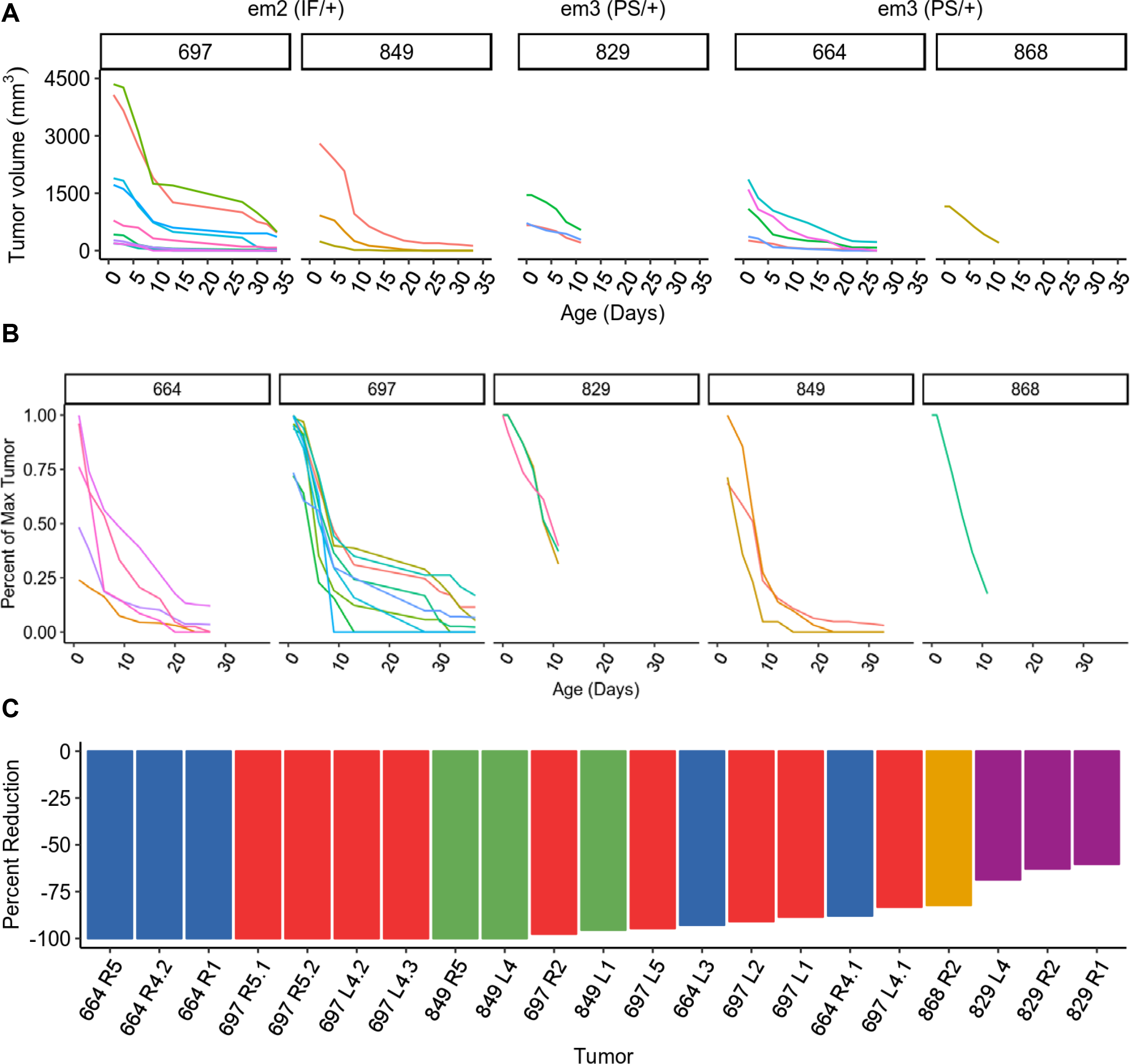
Estrogen depletion via ovariectomy results in rapid tumor regression in *Nf1*-deficient tumors. A) Tumor growth curves after overiectomies were performed (Day 0). B) Percent tumor loss of the largest tumor from each *Nf1* rat. C) The percent reduction in tumor size for all tumors from 3 *Nf1* rat lines is shown. Distinct colors are used to represent individual rats and the mammary pad location of each tumor is notated. Note that several rats had multiple measurable tumors within an individual mammary pad.

**Supplementary Table 1:**
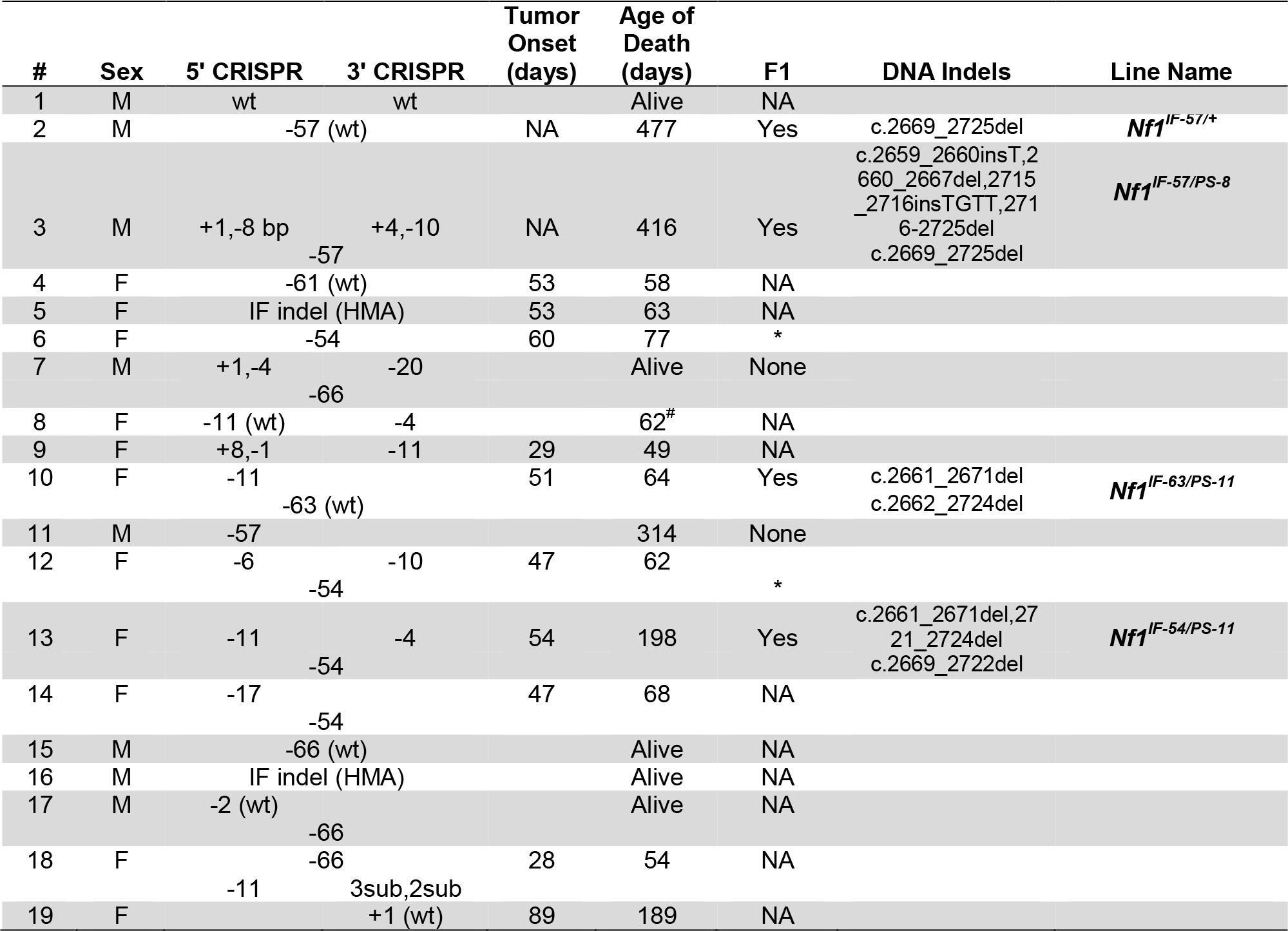
Summary of G_0_ animals obtained from CRISPR-Cas9 injections. Summary of indels resulting from CRISPR-Cas9 injections including tumor onset and survival. The most common base pair alterations at the 5’ and 3’ CRISPR sites are listed (additional indels that were identified in each G_0_ animal are listed in Supplemental Table 2). Mutant alleles that resulted in large deletions between 5’ and 3’ CRISPR target regions are listed in the middle of the 5’ and 3’ CRISPR columns; “wt” refers to the wildtype allele being observed by sequencing. IF indel (HMA) indicates that indels were identified by HMA but not sequence verified. All females were euthanized due to tumor burden except for #8 that was taken for control tissue (notated with #). DNA indel notations for non-founder animals can be found in Supplementary Table 2. * Euthanized when pregnant due to tumor burden. None = bred but no progeny.

**Supplementary Table 2:**
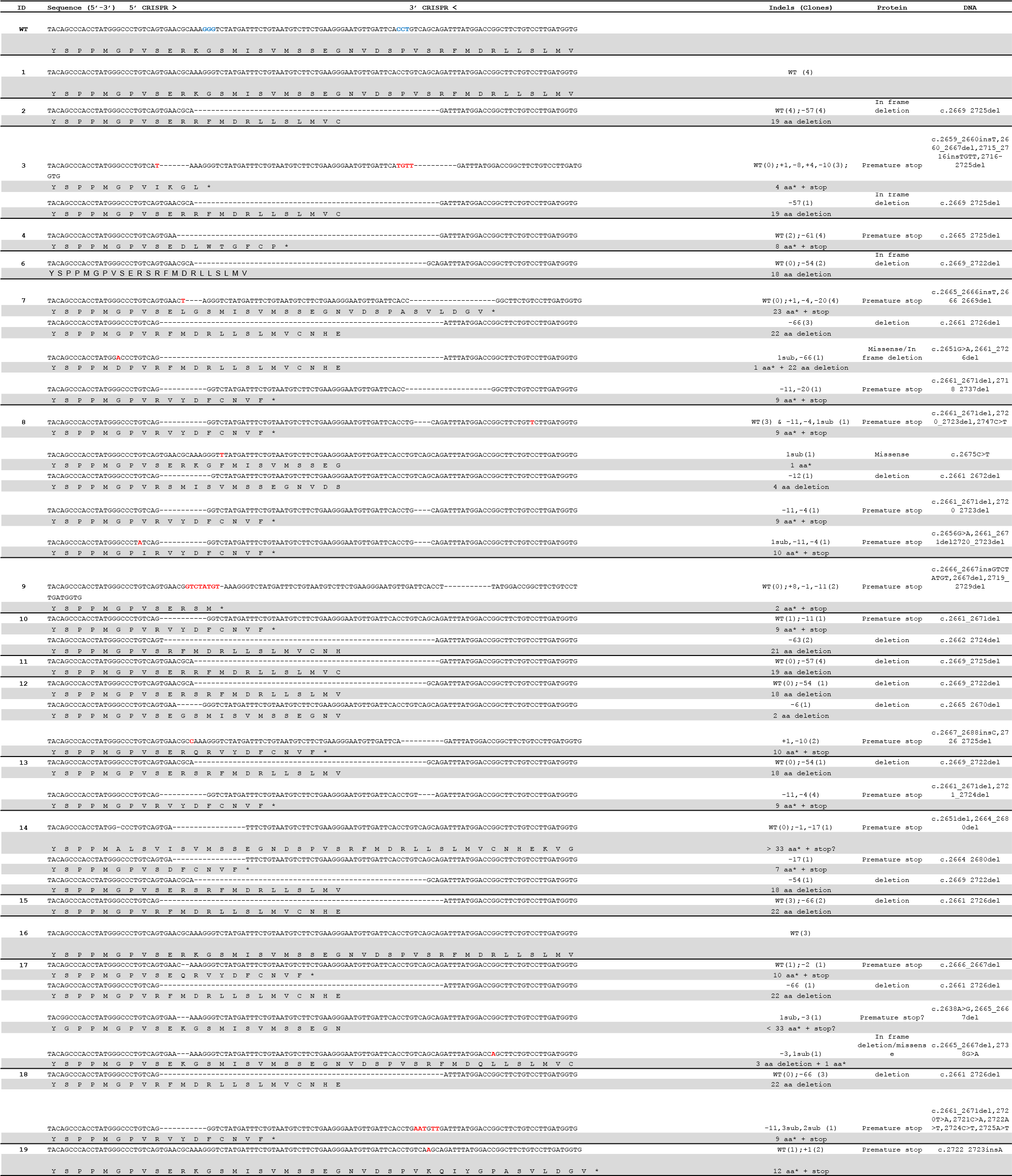
Detailed sequence analysis of *Nf1* G_0_ rats. 34 mutant alleles, including 25 unique mutant alleles, were identified by Sanger sequencing. Sequences for all alleles identified in G0 animals are shown.

**Supplementary Table 3:** Histopathology of *rNf1* animals.

| G0/F0 Animals |  |  |  |  |  |  |
| --- | --- | --- | --- | --- | --- | --- |
| Tumor/Slide |  |  |  |  |  |  |
| Animal | Sex | Genotype | Age (days) | No | Tissue/Mammary Pad Description | Pathology |
| 2 | M |  | 477 | 1 | R1 Mammary Fat Pad | mammary adenocarcinoma - acinar and mucinous |
|  |  |  |  | 2 | L4 Mammary Fat Pad | mammary pad - galactorrhea |
|  |  |  |  | 3 | R2 Mammary Fat Pad | mammary adenocarcinoma - acinar and ductular |
|  |  |  |  | 4 | L1 Mammary Fat Pad | two mammary adenocarcinomas 1-acinar and ductular 2-secretory acinar |
|  |  |  |  | 5 | L4 Mammary Fat Pad | mammary pad - galactorrhea |
|  |  |  |  | 6 | R1 Mammary Fat Pad | mammary pad - galactorrhea |
| 3 | M |  | 416 | 1 | L1 Mammary Fat Pad | mammary adenoma - cystic |
|  |  |  |  | 2 | Right flank | fibrous histiocytic sarcoma |
|  |  |  |  | 3 | L1 Mammary Fat Pad | mammary adenocarcinoma with two small intraductal tumors |
|  |  |  |  | 4 | L4 Mammary Fat Pad | mammary adenoma - cystic |
| 4 | F |  | 58 | 1 | R1 Mammary Fat Pad | mammary adenocarcinoma - acinar |
|  |  |  |  | 2 | L4 Mammary Fat Pad | mammary adenocarcinoma - acinar and solid |
|  |  |  |  | 3 | R4 Mammary Fat Pad | mammary adenocarcinoma - acinar and solid |
|  |  |  |  | 4 | L5 Mammary Fat Pad | mammary adenocarcinoma - acinar and solid |
|  |  |  |  | 5 | L4 Mammary Fat Pad | mammary adenocarcinoma - acinar and solid |
| 6 | F |  | 77 | 1 | R2 Mammary Fat Pad | mammary adenocarcinoma - acinar |
|  |  |  |  | 2 | R5 Mammary Fat Pad | mammary adenocarcinoma - acinar |
|  |  |  |  | 3 | L2 Mammary Fat Pad | mammary adenocarcinoma - acinar |
|  |  |  |  | 4 | L4 Mammary Fat Pad | mammary hyperplasia, mammary adenocarcinoma - acinar and ductular |
|  |  |  |  | 5 | R5 Mammary Fat Pad | mammary hyperplasia; mammary adenocarcinoma - acinar, ductular and cystic |
|  |  |  |  | 6 | L5 Mammary Fat Pad | mild mammay hyperplasia; mammary adenocarcinoma - cystic and papillary |
|  |  |  |  | 8 | R4 Mammary Fat Pad | mammary adenocarcinoma - ductular |
|  |  |  |  | N1 | L4-5 Mammary Pad | moderately severe acinar hyperplasia |
| 10 | F |  | 64 | 1 | R4 Mammary Fat Pad | mammary adenocarcinoma - acinar and solid |
|  |  |  |  | 2 | R5 Mammary Fat Pad | lactating mammary gland with small acinar adenocarcinoma |
|  |  |  |  | 3 | R5 Mammary Fat Pad | lactating mammary with adenocarcinoma - acinar and cystic |
|  |  |  |  | 4 | L4 Mammary Fat Pad | mammary adenocarcinoma - acinar and solid |
|  |  |  |  | 5 | L5 Mammary Fat Pad | mammary adenocarcinoma - acinar and solid |
|  |  |  |  | 6 | L5 Mammary Fat Pad | mammary adenocarcinoma - acinar |
| 13 | F |  | 198 | 1 | R4 Mammary Fat Pad | mammary adenocarcinoma - acinar and solid |
|  |  |  |  | 2 | L5 Mammary Fat Pad | 2 mammary adenocarcinomas - acinar and ductular 2) acinar |
|  |  |  |  | 3 | L3 Mammary Fat Pad | mammary adenocarcinoma - acinar and ductular |
|  |  |  |  | 4 | Right Brachial Lymph Node | large lymph node with hyperplasia |
|  |  |  |  | 5 | Left Brachial Lymph Node | large lymph node with hyperplasia |
|  |  |  |  | 6 | Aortic Lymph Node | lymph node |
|  |  |  |  | 7 | L4 Mammary Fat Pad | lymph node |
| rNF1-2 F1 animals |  |  |  |  |  |  |
| Tumor/Slide |  |  |  |  |  |  |
| Folder | Sex | Genotype | Age (days) | No | Tissue/Mammary Pad Description | Pathology |
| 303 | F |  | 62 | 1 | R4 Mammary Fat Pad | mammary adenocarcinoma - acinar and solid |
|  |  |  |  | 2 | R3 Mammary Fat Pad | mammary adenocarcinoma - solid, ductular with stroma |
|  |  |  |  | 7 | L4 Mammary Fat Pad | slight acinar hyperplasia |
|  |  |  |  | N1 | L4 normal mammary tissue | mild acinar hyperplasia |
| 304 | F |  | 89 | 1 | R4 Mammary Fat Pad | mammary adenocarcinoma - acinar and cystic |
|  |  |  |  | 2 | L5 Mammary Fat Pad | mammary adenocarcinoma - solid, acinar and cystic |
|  |  |  |  | 3 | L2 Mammary Fat Pad | mammary adenocarcinoma - solid and cystic |
|  |  |  |  | 4 | L1 Mammary Fat Pad | mammary adenocarcinoma - solid and cystic |
| 366 | F |  | 98 | 1 | L4 Mammary Fat Pad - 1 | mammary adenocarcinoma - acinar and ductular |
|  |  |  |  | 2 | L4 Mammary Fat Pad - 2 | mammary adenocarcinoma - acinar and ductular |
|  |  |  |  | 3 | R2 Mammary Fat Pad | mammary adenocarcinoma - acinar and ductular |
|  |  |  |  | 4 | R4 Mammary Fat Pad -1 | mammary adenocarcinoma - acinar and ductular |
|  |  |  |  | 5 | R4 Mammary Fat Pad - 2 | mammary adenocarcinoma - cystic and ductular |
|  |  |  |  | 6 | L2 Mammary Fat Pad | mammary adenoma |
| rNF1-3 F1 animals |  |  |  |  |  |  |
| Tumor/Slide |  |  |  |  |  |  |
| Folder | Sex | Genotype | Age (days) | No | Tissue/Mammary Pad Description | Pathology |
| 357 | F |  | 84 | 1 | R5 Mammary Fat Pad | mammary adenocarcinoma - mostly solid with some invasion |
|  |  |  |  | 2 | R1 Mammary Fat Pad | mammary adenocarcinoma - acinar |
|  |  |  |  | 3 | L4 Mammary Fat Pad - 2 | mammary adenocarcinoma - mostly solid with hemorrhage |
|  |  |  |  | 4 | L4 Mammary Fat Pad - 1 | mammary adenocarcinoma - solid |
|  |  |  |  | 9 | L2 Mammary Fat Pad | mammary adenocarcinoma - acinar and solid |
|  |  |  |  | 10 | R4 Mammary Fat Pad | mammary adenocarcinoma - acinar and solid |
|  |  |  |  | N1 | R3/R4 Normal Mammary Tissue | small mammary intraductal solid adenocarcinoma |
| 413 | F |  | 105 | 1 | L2 Mammary Fat Pad | mammary adenocarcinoma - acinar |
|  |  |  |  | 2 | L4 Mammary Fat Pad - 1 | mammary adenocarcinoma - acinar and ductular |
|  |  |  |  | 3 | L3 Mammary Fat Pad | mammary adenocarcinoma - acinar and solid |
|  |  |  |  | 4 | L4 Mammary Fat Pad - 2 | mammary adenocarcinoma - acinar and solid |
|  |  |  |  | 5 | L4 Mammary Fat Pad - 3 | 6 separate mammary tumors, 4 acinar, one adenoma and one with edema |
|  |  |  |  | 6 | L5 Mammary Fat Pad | solid, acinar, ductular and cystic |
|  |  |  |  | 7 | R2 Mammary Fat Pad | 5 small mammary acinar tumors in an abscess. |
|  |  |  |  | 8 | R5 Mammary Fat Pad | mammary adenocarcinoma - ductular, solid and acinar |
|  |  |  |  | 9 | R4 Mammary Fat Pad | mammary adenocarcioma - acinar, ductular, cystic |

| rNF1-10 F1 animals |  |  |  |  |  |  |
| --- | --- | --- | --- | --- | --- | --- |
| Folder | Sex | Genotype | Age (days) | Tumor/Slide |  | Pathology |
|  |  |  |  | No | Tissue/Mammary Pad Description |  |
| 246 | F |  | 96 | 1 | R2 Mammary Fat Pad | mammary adenocarcinoma - ductular and solid |
|  |  |  |  | 2 | L5 Mammary Fat Pad - 1 | mammary adenocarcinoma - cystic, papillary, solid, and ductal |
|  |  |  |  | 5 | L5 Mammary Fat Pad - 2 | mammary adenocarcinoma - ductular, acinar and cystic |
|  |  |  |  | N1 | R4 Normal Mammary Tissue | mild acinar hyperplasia |
| 252 | F |  | 183 | 1 | R4 Mammary Fat Pad | normal mammary gland |
|  |  |  |  | 2 | L5 Mammary Fat Pad | mammary adenocarcinoma - acinar |
|  |  |  |  | 3 | L4 Mammary Fat Pad | focal hyperplasia and mammary adenoma |
|  |  |  |  | 7 | L2 Mammary Fat Pad | mammary adenocarcinoma - solid |
|  |  |  |  | N1 | L2 Mammary Fat Pad | normal mammary |
| rNF1-13 F1 animals |  |  |  |  |  |  |
| Folder | Sex | Genotype | Age (days) | Tumor/Slide |  | Pathology |
|  |  |  |  | No | Tissue/Mammary Pad Description |  |
| 262 | F |  | 92 | 1 | R2 Mammary Fat Pad | mammary adenocarcinoma - solid, ductular with stroma |
|  |  |  |  | N1 | Normal mammary tissue | normal mammary tissue |
| 391 | F |  | 89 | 1 | R4 Mammary Fat Pad - 1 | mammary adenocarcinoma - ductular |
|  |  |  |  | 2 | R4 Mammary Fat Pad - 2 | mammary adenocarcinoma - ductular and solid |
|  |  |  |  | 3 | R4 Mammary Fat Pad - 3 | mammary adenocarcinoma - ductular |
|  |  |  |  | 4 | L4 Mammary Fat Pad | 3 small mammary adenocarcinomas - acinar |
|  |  |  |  | 5 | L2 Mammary Fat Pad | mammary adenocarcinoma - acinar and solid |
|  |  |  |  | N1 | R2 Normal Mammary | normal mammary gland |
| 397 | F |  | 118 | 1 | L2 Mammary Fat Pad | mammary adenocarcinoma - solid, acinar and cystic |
|  |  |  |  | 2 | L4 Mammary Fat Pad - 2 | mammary adenocarcinoma - acinar and solid |
|  |  |  |  | 3 | L5 Mammary Fat Pad | mammary adenocarcinoma - solid |
|  |  |  |  | 4 | R5 Mammary Fat Pad | mammary adenocarcinoma - solid and acinar |
|  |  |  |  | 5 | L4 Mammary Fat Pad - 1 | mammary adenocarcinoma - ductular |
|  |  |  |  | N1 | R4 Normal Mammary | mildly hyperplastic mammary gland |

